# Workflow enhancement of TurboID-mediated proximity labeling for SPY signaling network mapping

**DOI:** 10.1101/2024.02.17.580820

**Authors:** TaraBryn S Grismer, Sumudu S Karundasa, Ruben Shrestha, Danbi Byun, Weimin Ni, Andres V Reyes, Shou-Ling Xu

## Abstract

TurboID-based proximity labeling coupled to mass spectrometry (PL-MS) has emerged as a powerful tool for mapping protein-protein interactions in both plant and animal systems. Despite advances in sensitivity, PL-MS studies can still suffer from false negatives, especially when dealing with low abundance bait proteins and their transient interactors. Protein-level enrichment for biotinylated proteins is well developed and popular, but direct detection of biotinylated proteins by peptide-level enrichment and the difference in results between direct and indirect detection remain underexplored. To address this gap, we compared and improved enrichment and data analysis methods using TurboID fused to SPY, a low-abundance O-fucose transferase, using an AAL-enriched SPY target library for cross-referencing. Our results showed that MyOne and M280 streptavidin beads significantly outperformed antibody beads for peptide-level enrichment, with M280 performing best. In addition, while a biotin concentration ≤ 50 µM is recommended for protein-level enrichment in plants, higher biotin concentrations can be used for peptide-level enrichment, allowing us to improve detection and data quality. FragPipe’s MSFragger protein identification and quantification software outperformed Maxquant and Protein Prospector for SPY interactome enrichment due to its superior detection of biotinylated peptides. Our improved washing protocols for protein-level enrichment mitigated bead collapse issues, improving data quality, and reducing experimental time. We found that the two enrichment methods provided complementary results and identified a total of 160 SPY-TurboID-enriched interactors, including 60 previously identified in the AAL-enriched SPY target list and 100 additional novel interactors. SILIA quantitative proteomics comparing WT and *spy-4* mutants showed that SPY affects the protein levels of some of the identified interactors, such as nucleoporin proteins. We expect that our improvement will extend beyond TurboID to benefit other PL systems and hold promise for broader applications in biological research.

## Introduction

Proximity labeling mass spectrometry (PL-MS) is a powerful technique that has been widely used to map protein-protein interactions (PPIs), subcellular proteomes, cell-type proteomes, and protein-RNA and protein-DNA interactions (1, 2). TurboID, a promiscuous engineered variant of biotin ligase with greatly enhanced activity, has recently been widely used in both animal and plant research fields (3). By fusing TurboID to a protein of interest or subcellular marker, or expressing it under a cell-specific promoter, TurboID facilitates the labeling of proximal proteins with biotin, followed by the isolation of these biotinylated proteins by affinity purification and mass spectrometric analysis. Because protein complexes don’t need to remain associated during the purification process, PL-MS bypasses several hurdles of traditional methods, such as IP-MS or cellular fractionation, and provides unprecedented sensitivity, marking a major advance in the study of complex biological interactions and subcellular proteomic analysis.

Despite the versatility, simplicity, and greatly improved sensitivity of PL-MS compared to traditional methods, PL-MS still presents some challenges. For low abundance baits and/or their weak/transient interactors, PL-MS can still produce false negatives. This is partly due to the large amount of background proteins that persist despite extensive washes. For example, even though the streptavidin magnetic beads have nearly the highest binding affinity and the wash conditions are harsh, we have observed that over 1900 background proteins (>15,000 peptides) are routinely detected in wild-type samples without TurboID (4), which can potentially obscuring the detection of true low-abundance candidates. Increasing the number of beads or input materials is often not an option, as the background is already high. On the other hand, increasing the concentration of the biotin is not an option for enrichment of biotinylated proteins at the protein level because high concentrations of biotin are not easily removed and compete with the bead binding (4, 5). Further optimization is needed to increase the sensitivity of PL-MS.

Many researchers have used protein-level enrichment of biotinylated proteins (3, 4, 5, 6), while studies focusing on peptide-level enrichment are relatively limited (7, 8). It is generally believed that the protein-level enrichment may provide higher sensitivity because any peptide identified from the captured protein is considered valid. In contrast, peptide-level enrichment must identify biotinylated peptides, which are present at much lower abundance compared to unmodified peptides, resulting in potentially lower detection sensitivity. However, enrichment of biotinylated peptides at peptide-level followed by MS analysis is considered direct detection (9) and has the advantage of eliminating potential false positives arising from the high background proteins from protein-level enrichment. In addition, the sample complexity of peptide-level enriched samples is expected to be significantly lower than protein-level enriched samples, which may allow the detection of low-abundance biotinylated prey peptides possible. A direct comparison between the two approaches is still lacking.

Here, we aim to increase the sensitivity of PL-MS in several steps. We chose SPY-TurboID lines as our experimental model, exploiting SPY, an Arabidopsis O-fucose transferase, known for its low abundance and thought to have weak transient interactions with numerous targets (10–12). We previously generated an Aleuria aurantia lectin (AAL)-enriched library of SPY-modified targets (12), which was used to cross-reference our results in this study. For peptide-level enrichment, we systematically tested different bead types, quantification methods, and biotin concentrations to increase detection sensitivity. In addition, we optimized the washing procedure to prevent bead collapse during protein-level enrichment, thereby improving overall sensitivity. We directly compared the results of the two methods, considering both the number of detected known SPY targets and the number of unique interactors. Our results show that peptide-level and protein-level enrichment provide complementary results. In total, we detected 60 known targets, but 100 novel interactors. We detected 10 members of the nucleoporin proteins as SPY targets, and our comprehensive quantitative analysis comparing the proteomes of WT and *spy-4* mutants shows that SPY affects the protein levels of several members.

## Results

### Comparison Workflow and Selected Lines

We established a workflow to compare and optimize the biotinylated protein enrichment at both the peptide and protein level and to compare data analysis software (Fig. 1A). The SPINDLY-TurboID-VENUS (SPY-TD) line, expressed under its native promoter in the *spy-3* mutant background, was used for testing (Fig. 1B). SPY functions as an O-fucosyltransferase that adds fucose to serine and threonine residues on a variety of nucleocytosolic acceptor substrates (11, 12). The SPY protein is localized to the nucleus and the cytosol (13). It exhibits relatively modest protein expression levels, with a median iBAQ of 22.48. This is 10-fold and 8-fold log2 difference compared to Rubisco (32.79) and Actin 2 (30.64), respectively (14). In a recent study, we used AAL chromatography to enrich SPY targets and identified 467 substrates from Arabidopsis seedlings and flower tissues (12). Given the large number of substrates and the relatively low abundance of SPY, the interaction between SPY and substrates is predicted to be weak and transient. We reasoned that this setup would allow us to evaluate the performance of biotinylated protein enrichment and make adjustments to improve efficiency.

**Figure 1.**
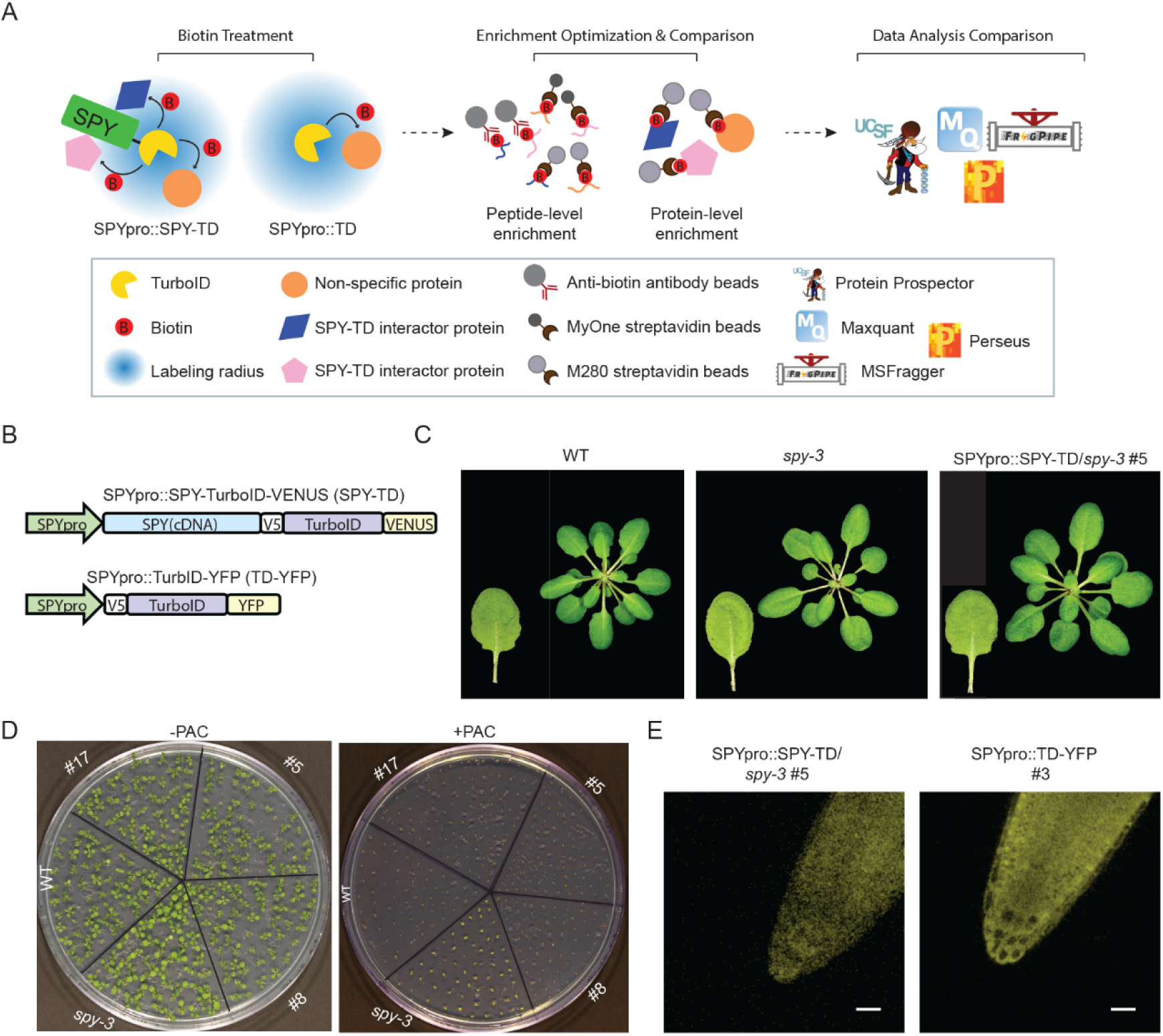
Schematic workflow for comparison and optimization of biotinylated protein enrichment and SPY-TD lines used for testing. (A) Comparison and optimization for both peptide- and protein-level enrichment. (B) Schematic constructs for bait SPY-TurboID-VENUS (SPY-TD) and control TurboID-YFP (TD-YFP) expressed under the native SPY promoter. (C) SPY-TD fusion protein complements the leaf serration phenotype of *spy-3*. (D) The germination on PAC phenotype of *spy-3* was rescued to WT level by the SPY-TD fusion. (E) Confocal images of SPY-TD-VENUS and TD-YFP show nuclear and cytosolic localization for both bait and control. scale= 25 µm.

To ensure that the SPY-TD fusion protein is fully functional, we first examined its complementation in the *spy-3* mutant. The *spy-3* mutant has a smooth leaf phenotype (Fig. 1C) and can germinate in the presence of paclobutrazol (PAC) (Fig. 1D) (15, 16). As expected, the SPY-TD lines completely rescued the leaf serration and germination on PAC phenotype of the *spy-3* mutant phenotype to the wild type (Fig. 1C and 1D). These complementation results demonstrate that the SPY-TD fusion protein is fully physiologically functional. As a control, TurboID fused to YFP (TD-YFP) was expressed under the same native SPY promoter (TD-YFP) and a similar expression level expression line to SPY-TD was selected. Confocal microscopy confirmed that both SPY-TD and TD-YFP are expressed both in the nucleus as well as in the cytosol (Fig. 1E).

We then performed a biotin treatment on the SPY-TD line (Supplementary Fig. S1), ranging from 0 to 1000 µM of biotin treatment, followed by a Western blot using α-streptavidin. Compared to the wild type, we observed three bands showing specific biotinylation in SPY-TD, including one band of the predicted bait fusion (SPY-TD) itself. However, the intensity of these bands did not show an obvious increase with increasing biotin concentration. This suggests that either the interaction between SPY and its interactors is very weak/transient, resulting in weak labeling, and/or that most of its substrates are of very low abundance and undetectable by α-streptavidin.

### M280 Streptavidin Beads Outperform Other Bead Types for Peptide-Level Enrichment

Using the SPY-TD line, we first started with biotinylated peptide enrichment, which allows direct detection of biotinylated proteins (9). We compared anti-biotin antibody agarose beads, MyOne streptavidin and M280 streptavidin beads (Fig. 2A). Ten-day-old Arabidopsis seedlings of the SPY-TD line were treated with biotin at a concentration of 50 µM for 1 h, which is commonly used for protein-level enrichment (4, 5). The proteins were then extracted and digested to peptides. The resulting approximately 4.5 mg of peptides were used as input for each enrichment (see Methods). Data analysis was performed using the Protein Prospector search engine, which allows the target decoy calculation for biotinylated peptides (17, 18).

**Figure 2.**
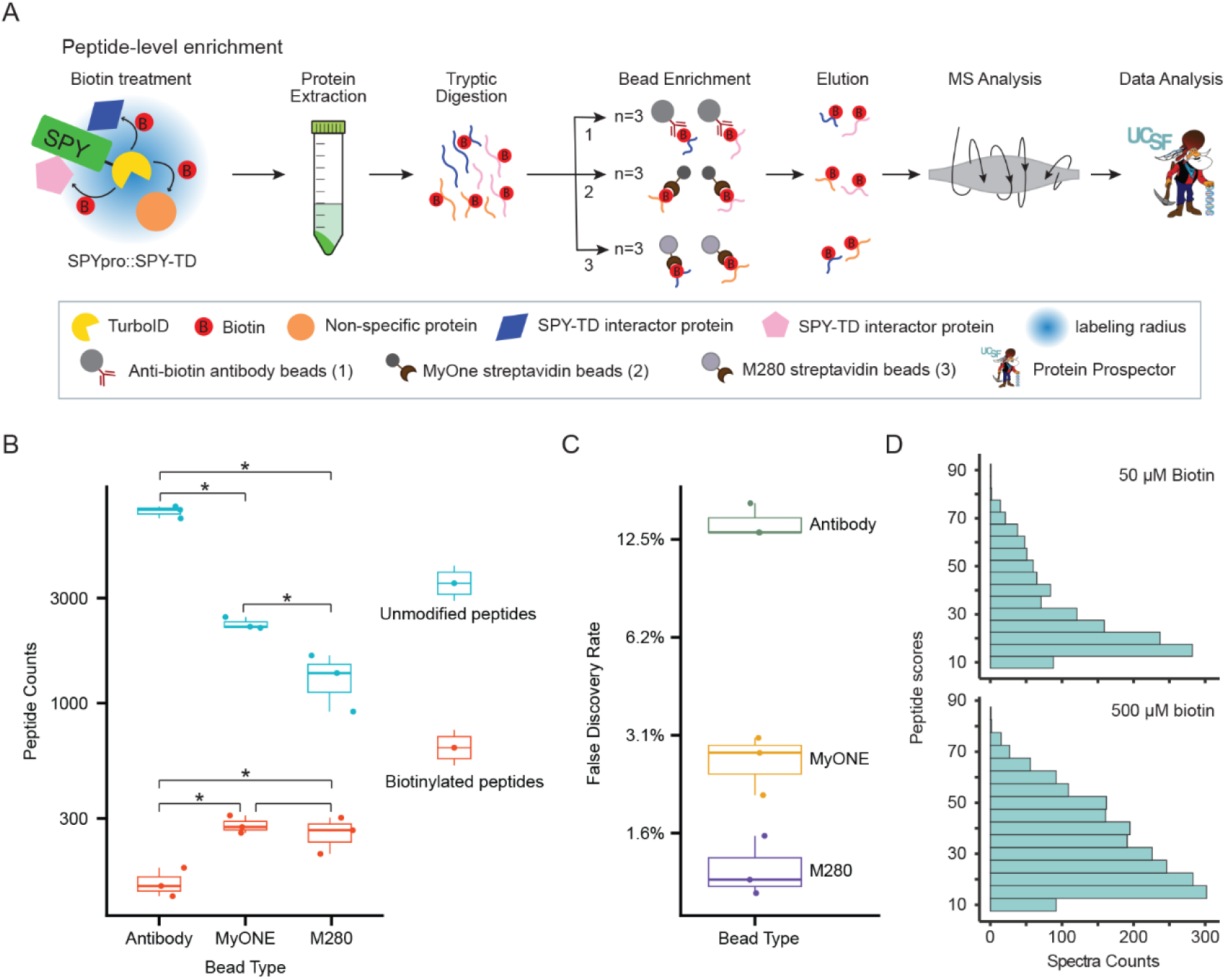
Peptide-level enrichment of biotinylated proteins shows better performance of M280 beads and higher biotin concentration. (A) Schematic workflow for peptide-level biotinylation enrichment comparing two parameters: bead type and biotin concentration. (B) M280 beads outperform anti-biotin antibody and MyOne beads in peptide-level enrichment. Modified and unmodified peptide counts identified by each bead type (in three biological replicates of SPY-TD lines treated with 50 µM biotin) were compared to check for statistical difference using a two-tailed t-test (*, p <0.05). (C) M280 beads have a lower false discovery rate (FDR) compared to the other two bead types. FDRs for each bead type were calculated as the percentage of biotinylated decoy peptides identified out of the total biotinylated peptides. (D) A higher concentration of biotin treatment (500 µM) results in improved identification with higher peptide scores and a greater abundance of biotinylated peptides than the 50 µM treatment.

We first tested antibody-based agarose bead enrichment (8), which allowed elution of biotinylated peptides in 0.15% TFA but limited extensive washing. Using a 1% false discovery rate (FDR) for peptides and a 5% FDR for protein identification, three replicates yielded an average of 153 biotinylated peptides and 7421 unmodified peptides (Fig. 2B, Supplementary Table 1), corresponding to 109 modified proteins and 2748 unmodified proteins. Biotinylated peptides were detected from endogenously biotinylated proteins (e.g. MCCA, BCCP2) and known SPY targets (e.g. NEDD1, PUM2). However, calculation of the FDR based solely on target decoy results for biotinylated peptides (17) showed an average FDR of 13.8% (Fig. 2C, Supplementary Table 1).

Next, we investigated streptavidin-based enrichment using the protocol optimized for click chemistry-based enrichment of biotinylated peptides (9). Streptavidin’s high affinity for biotin (Kd = 10^-14^) requires a strong elution (0.2% TFA, 0.1% FA, and 80% ACN at 75°C) but allows for more washing to remove nonspecific interactors. MyOne (1-μm) and M280 (2.8-μm) magnetic streptavidin beads, commonly used in many laboratories for protein-level enrichment of biotinylated proteins, were tested (3–5).

MyOne yielded an average of 280 biotinylated peptides and 2293 unmodified peptides, while M280 yielded 258 biotinylated peptides and 1310 unmodified peptides (Fig. 2B). Positive controls such as NEDD and PUM2 were consistently detected. The FDR calculated for biotinylated peptides was 2.5% for MyOne and 1.2% for M280 (Fig. 2C, Supplementary Table 1).

Among the three types of beads, antibody agarose beads showed poor performance with lower identification of biotinylated peptides, higher background noise, and higher FDR. While higher background is generally manageable for identification with modern mass spectrometers, antibody enrichment resulted in a higher FDR for modified peptides due to a greater number of background unmodified peptides. In contrast, MyOne and M280 beads showed better results in terms of number of biotinylated peptides identified, background noise, and FDR. In particular, M280 showed excellent results for low background noise and FDR. Therefore, M280 beads were selected for further experiments.

### High Concentration Biotin Treatment Provides Better Peptide-Level Enrichment Results

We next investigated methods to improve the detection and quantification of the SPY interactome. Despite successful detection of a number of known targets, we observed a prevalence of unmodified peptides (Fig. 2B) and generally low scores for modified peptides. To improve performance, we considered several optimization measures, including increasing the concentration of acetonitrile in the enrichment process, increasing the concentration of biotin for treatment, and testing different search engines.

Given the increased peptide hydrophobicity due to the biotin modification, we tested 5%, 10% and 15% acetonitrile concentrations (Supplementary Fig. S2). The results showed that 15% gave slightly better results—lower FDR and higher percentage of biotinylated peptides – without affecting streptavidin binding (Supplementary Table 2). Therefore, we proceeded with 15% ACN for peptide-level enrichment.

Next, we investigated a higher concentration of biotin treatment. In plants, the recommended biotin concentration for protein-level enrichment is typically around 50 µM. Higher concentrations of biotin are not recommended due to the challenge of biotin removal by washing. Even at 50 µM, biotin must be removed using a desalting column, such as the PD-10 column, to prevent free biotin from competing with the streptavidin binding (1, 4, 5). However, these columns have a limited ability to remove free biotin, so in general significantly more streptavidin beads must be used, resulting in an increase in background noise. For peptide-level enrichment, however, free biotin can be effectively removed during the protein precipitation.

We chose a higher concentration of 500 µM biotin for the enrichment and compared it with 50 µM enrichment using a similar input (∼10 mg peptides) of SPY-TD samples. The histograms in Fig. 2D show more detected biotinylated peptides by spectra counting from a total of three replicates, and importantly, their peptide scores are significantly higher than those from 50 µM (Supplementary Table 2). This result shows that the high biotin treatment provides much more reliable results for peptide-level enrichment.

### MSFragger Provides Better Results Than Maxquant for Peptide-Level Enrichment

Next, we performed a quantitative analysis to compare SPY-TD with the TD-YFP control treated with 500 µM biotin to identify SPY-specific interactors. For this comparison, we used both Maxquant (using the Andromeda search engine) (19) and FragPipe’s MSFragger search engine (20) (Fig. 3A), free publicly available software. Both Maxquant and MSFragger facilitate label-free quantification (LFQ) by extracting peptide intensities, supporting match between runs, and using the statistical tool Perseus (21). It’s worth noting that the Protein Prospector lacks similar features.

**Figure 3.**
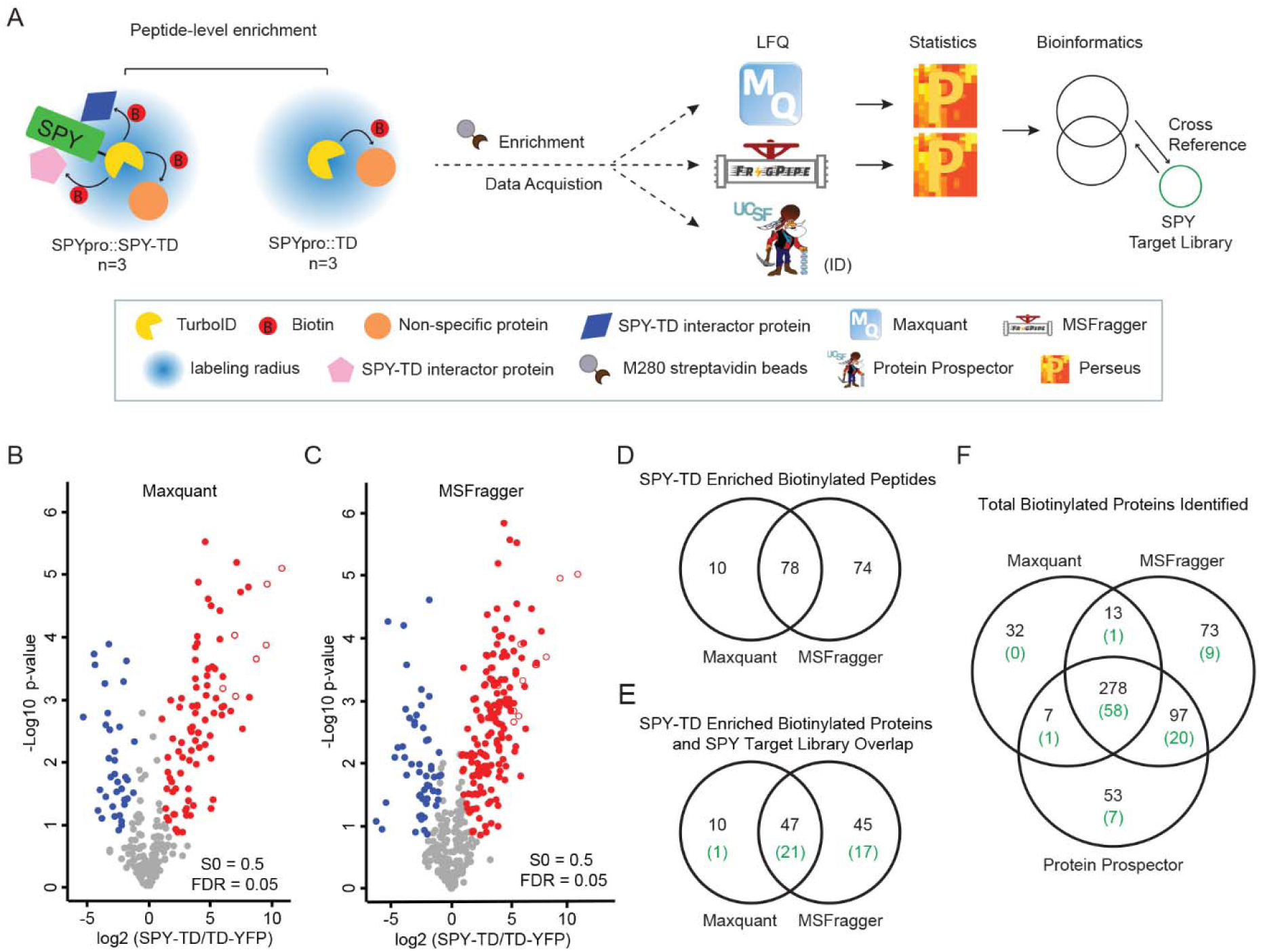
MSFragger software shows better performance than Maxquant for peptide-level enrichment detection and quantification of biotinylated proteins. (A) Schematic workflow for software comparison to determine the SPY-TD specific interactome. SPY-TD and control were enriched at the peptide-level followed by label-free quantification (LFQ) using Maxquant and MSFragger. Enriched proteins are compared to the SPY target library for comparison. (B-C) Quantitative analysis of SPY-TD peptide-level enrichment using Maxquant (B) and MSFragger (C). SPY-TD enriched peptides are shown in red; control enriched peptides are shown in blue. Biotinylated peptides detected for the SPY bait itself are shown as empty circles. (D) Venny plot comparing SPY-TD enriched biotinylated peptides between Maxquant and MSFragger showing more enriched SPY-TD biotinylated peptides from MSFragger analysis. (E) Venny plot comparing SPY-TD enriched biotinylated peptides grouped to proteins between Maxquant and MSFragger. Both shared and unique biotinylated proteins are cross-referenced to the SPY target library, and the overlap number is shown in parentheses, colored in green. (F) Comparison of the total number of identified biotinylated proteins between three software programs, prior to any filtering or statistical tests. Detected known targets that overlapped with each catalog list are shown in the parentheses, colored in green.

Using LFQ and Perseus on biotinylated peptides from the Maxquant input, our quantitative analysis revealed 88 modified peptides corresponding to 57 proteins significantly enriched in the SPY-TD samples, including SPY itself (empty circles) (Fig. 3B) (Supplementary Table 3). In contrast, LFQ and Perseus analysis using MSFragger on the same datasets revealed 152 modified peptides corresponding to 92 proteins enriched in SPY-TD samples (Fig. 3C) (Supplementary Table 3). Notably, there was an overlap of 78 biotinylated peptides (47 proteins) between the results of the two software, and unique identifications by both are shown in Fig. 3D (peptide overlap) and Fig. 3E (protein overlap).

To further assess enrichment, we compared our enriched proteins with the AAL-enriched SPY target library (12) (Fig. 3E). Interestingly, 47 proteins were found to be enriched by both Maxquant and MSFragger, 21 of which were supported by the SPY target library. MSFragger showed 45 proteins uniquely enriched by SPY-TD, 17 of which were listed in the SPY target library. In contrast, Maxquant showed only 1 out of 10 uniquely enriched proteins listed in the SPY target library. Based on these results, we conclude that MSFragger is more effective for quantifying biotinylated peptide enrichment using LFQ.

To investigate whether the discrepancy between search engines was due to differences in identification rates, which could affect downstream quantification, we compared the list of total biotinylated peptides (Supplementary Fig. S3A, Supplementary Table 4) and proteins (Fig. 3F) identified from six runs, including both SPY-TD and control runs, between Maxquant and MSFragger, and Protein Prospector. Specifically, MSFragger identifies the most biotinylated proteins (753 peptides/ 461 proteins), Protein Prospector identifies 674 peptides/435 proteins, and Maxquant identifies the fewest biotinylated proteins (497 peptides/327 proteins). In total, 426 peptides/278 biotinylated proteins were detected by all three software programs (Fig. 3F, Supplementary Fig. S3A). MSFragger and Protein Prospector identified an additional 97 common proteins, 20 of which matched the SPY target library (Fig. 3F). These results suggest that the superior performance of MSFragger is mainly due to the higher identification rates for the biotinylated peptides of the MSFragger search engine. Notably, we observed that certain highly scored biotinylated peptides, such as a peptide from TCP22 (Supplementary Fig. S3B), which were enriched in SPY-TD samples by Skyline quantification (Supplementary Fig. S3C), were identified by Protein Prospector but were missing in MSFragger. This suggests that MSFragger, although the best performer of the three, may still be missing some identifications of biotinylated peptides.

We observed that signature ions such as the derivative ion at m/z 310.158 due to ammonia loss from the immonium ion of biotinylated lysine (ImKBio) are robustly detected in our HCD MS2 spectra of biotinylated peptides (Supplementary Fig. S3B), consistent with previous findings (22, 23), while other signature ions such as m/z 227.085 and 327.185 are less observed, with the 327.185 peak being the least observed.

### Optimized Wash Procedures for Improved Protein-Level Biotinylation Enrichment and Improved Quantification

During protein-level enrichment, we frequently observed bead collapse (beads adhering to the tube walls and not completely pulled to the side of the tube by the magnetic force), potentially resulting in sample loss (Fig. 4B-C). As a result, we conducted a detailed review of the protocol (3, 4). Prior to elution by tryptic digestion, streptavidin beads are extensively washed with extraction buffer containing detergent (SDS), high salt (1M KCl), high pH (0.1M Na_2_CO_3_), and chaotropic denaturants (2M urea). We observed that bead collapse became more pronounced after the 1M Na_2_CO_3_ and urea washes (Fig. 4B). Therefore, in our modified washes, where we replaced the Na_2_CO_3_ wash with an extraction buffer wash followed by urea washes, we observed a significant reduction in bead collapse (Fig. 4A and C, Supplementary Fig. S4).

**Figure 4.**
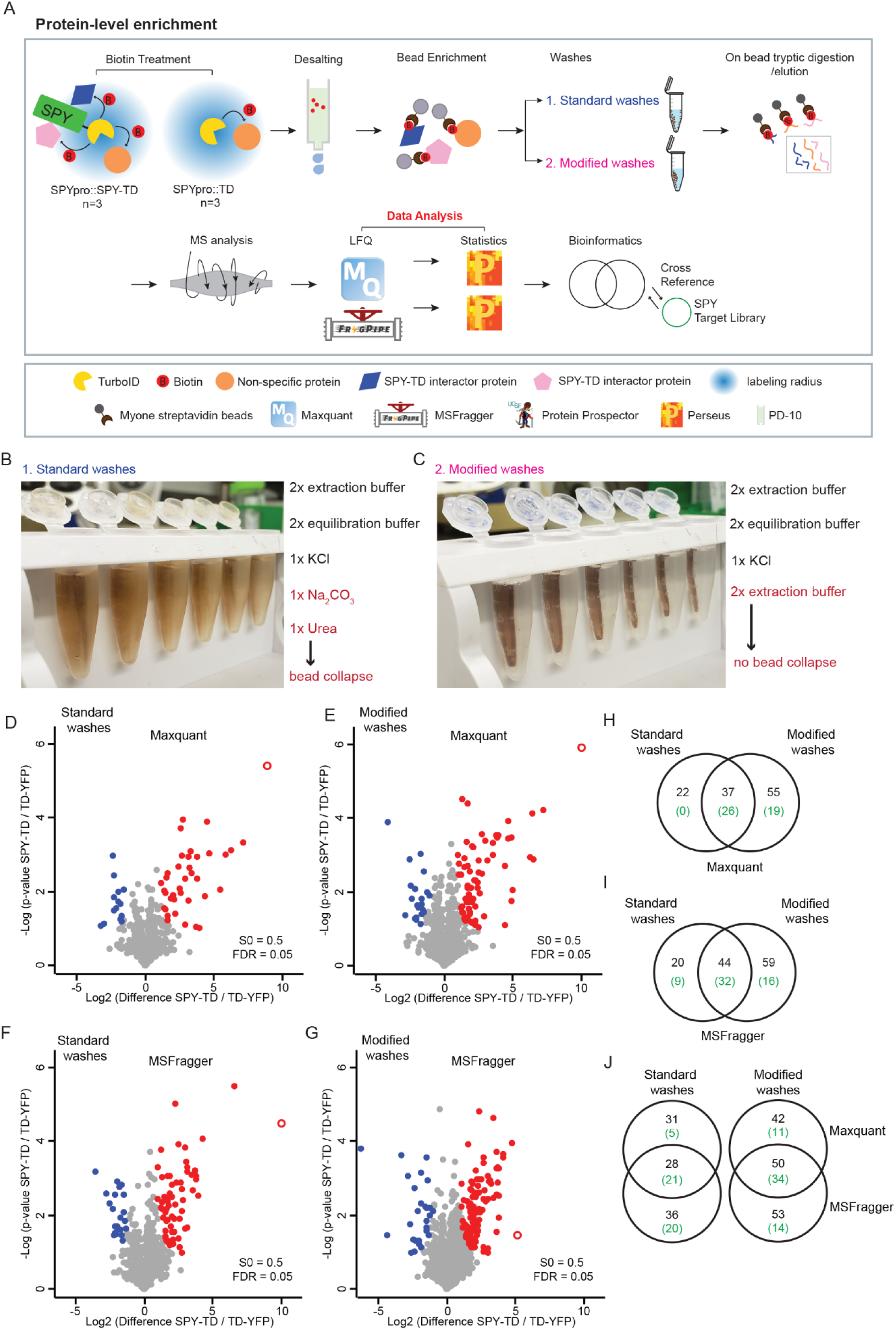
Modified washes outperform standard washes for protein-level enrichment of biotinylated proteins. (A) Schematic workflow to optimize washes for protein-level enrichment of biotinylated proteins. (B-C) Bead collapse observed in standard washes (B) but not in the modified washes (C). Image shows bead collapse at the urea wash step after the Na_2_CO_3_ wash. (D-G) Volcano plots of SPY-TD enriched proteins comparing standard and modified washes using Maxquant and MSFragger. Red and blue dots represent proteins showing enrichment in SPY-TD samples and TD-YFP samples, respectively. The bait SPY is shown as an empty circle. (H-I) Both Maxquant and MSFragger show that modified washes provide better enrichment results as more enriched proteins are identified. The overlap with the SPY target library is shown in parentheses, colored green. (J) Maxquant and MSFragger each enrich unique proteins in both standard washes and modified washes. Overlap with the SPY target library is shown in parentheses, colored green.

To test whether the modified washes improved the enrichment quantification, we compared the MS results using LFQ (S0=0.5, FDR=0.05) and analyzed using both Maxquant (Fig. 4A, D-E) and MSFragger (Fig. 4A, F-G). We observed that the modified wash yielded more enriched proteins than standard washes using either Maxquant (92 vs 59) or MSFragger (103 vs 64) (Fig. 4H-I), and more known SPY targets overall using Maxquant (45 vs 26) and using MSFragger (48 vs 41) (Fig. 4H-I).

To check which software produced better data, we also compared Maxquant with MSFragger (Fig. 4J). Overall, MSFragger enriched more SPY-TD proteins than Maxquant. In the standard washes, 28 interactors are enriched in both; MSFragger appears to enrich a higher percentage of SPY known targets than Maxquant for uniquely enriched proteins (20/36 vs 5/31). However, in the modified washes, it appears that both MSFragger and Maxquant quantified unique SPY-TD enriched proteins with a similar percentage (14/53 vs 11/42), but MSFragger has a slightly higher number of uniquely enriched interactors. We note the differences in quantification of the SPY bait protein (red empty circle) in modified washes, which appears highly enriched in Maxquant data analysis (log2 fold difference 10.018, −log10 p-value 5.893), but shows less enrichment in MSFragger (log2 fold difference 5.135, −log10 p-value 1.457) (Fig. 4E and G), while in standard washes, MSFragger shows similar quantification results of SPY as Maxquant (Fig. 4D and F).

### Peptide-Level Biotinylation Enrichment Provides Complementary Results to Protein-Level Enrichment

We then selected the more optimal MSFragger-analyzed results from each method and compared the results obtained by peptide-level enrichment and protein-level enrichment. Protein-level enrichment revealed 103 proteins enriched by SPY-TD, 48 of which were matched to the SPY target library. In contrast, peptide-level enrichment revealed 92 proteins enriched by SPY-TD, 38 of which were matched to SPY targets (Fig. 5A). Notably, 35 proteins were common to both methods, with peptide-level enrichment yielding 57 unique proteins and protein-level enrichment yielding 68 unique proteins. 12 out of these 57 unique proteins from the peptide-level enrichment matched the SPY target library, and 22 out of the 68 unique proteins from the protein-level enrichment matched the SPY target library (Supplementary Table 5). This suggests that protein-level enrichment provides better overall results, while peptide-level enrichment provides complementary results. On the other hand, peptide-level enrichment provides compelling direct evidence with its direct biotinylation identification.

**Figure 5.**
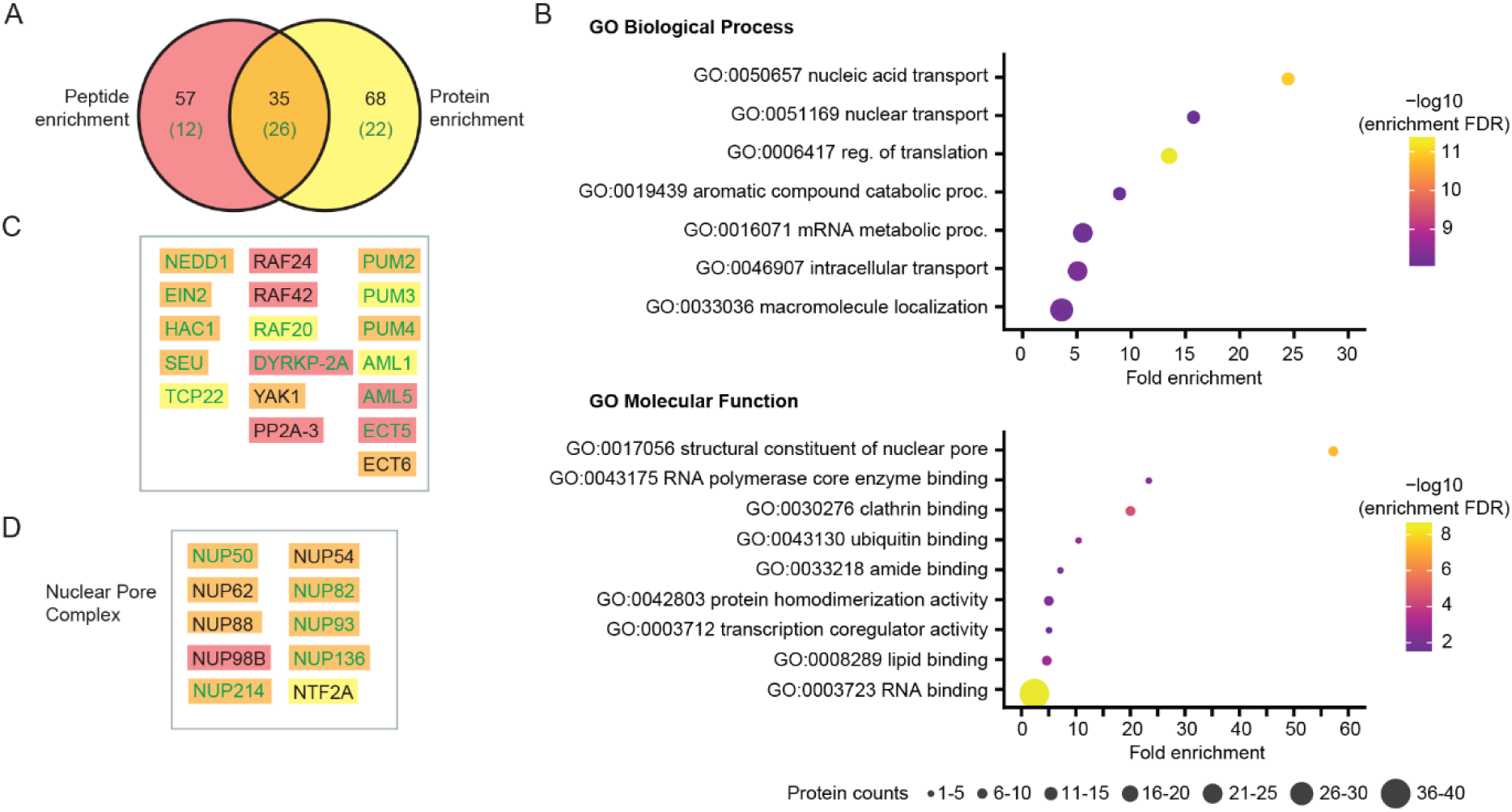
Biotinylation enrichment at the peptide and protein level provides complementary enrichment results and adds information to the SPY target library. (A) Venn diagram of SPY-TD enriched proteins at the peptide and protein level showing the overlapping (colored in orange) but also unique interactors (peptide-unique colored in red, protein-unique colored in yellow). Overlap with the SPY target library is shown in parentheses, colored green. (B) ShinyGO analysis of SPY-TD interactors reveals enriched biological processes and molecular functions. (C) Examples of proteins with the same color theme in (A). PL-MS of SPY-TD identifies known and novel interactors, including additional transcription factors and phosphatase/kinases. (D) PL-MS of SPY-TD significantly enriches the nuclear pore protein family.

We then performed GO analysis on these identified proteins (Fig. 5B) for biological process and molecular function. Not surprisingly, proteins involved in several regulatory processes are enriched, including macromolecular localization, intracellular transport and mRNA metabolic processes. For molecular function, proteins show a significant enrichment in RNA binding.

Known components previously identified as SPY targets, such as NEDD1, EIN2, PUM2, HAC1, SEU, AML1, TCP22, were detected by PL-MS (Fig. 5C). More importantly, more family members of known targets are identified by PL-MS; for example, in addition to RAF20, we now detect RAF24 and RAF42; in addition to the known ECT5, we now detect ECT6. And, in addition to previously identified members of the nuclear pore complex, we now identify 5 new proteins involved (Fig. 5D).

Both the protein- and the peptide-level enrichment surpass our previous attempts using IP-MS with SPY-GFP bait (13) (Supplementary Fig. S5). Our IP-MS identified only four significant enriched protein groups, including the SPY bait and known target TCP14 (24). These results highlight the improved capabilities of PL-MS in identifying transient and weak interactors.

### SPY Affects Nucleoporin Protein Levels

Since several nucleoporin proteins were identified either as SPY targets or in the SPY interactome data of this study, and the structural constituent of the nuclear pore is significantly enriched by GO analysis (Fig. 5B and D), we asked whether SPY affects the protein levels of nucleoporin. We performed stable isotope labeling analysis using ^15^N labeling (18). Four replicates of the SILIA quantitative analysis were performed and processed using Protein Prospector for quantification (Fig. 6A). Proteins quantified in three out of the four replicates were retained and filtered (Supplementary Table 6). Of the final 4910 proteins quantified, we found that at least 3 nucleoporin proteins are significantly lower in the *spy-4* mutants, including NUP50, NUP82, and NUP136 (NUP1) with more than 2-fold reduction, with NUP50 showing the largest reduction (Fig. 6B and C). NUP 155, NUP93, and NUP 85 also show a significant reduction compared to the tubulin and actin control, although the reduction is milder (Fig. 6C).

**Figure 6.**
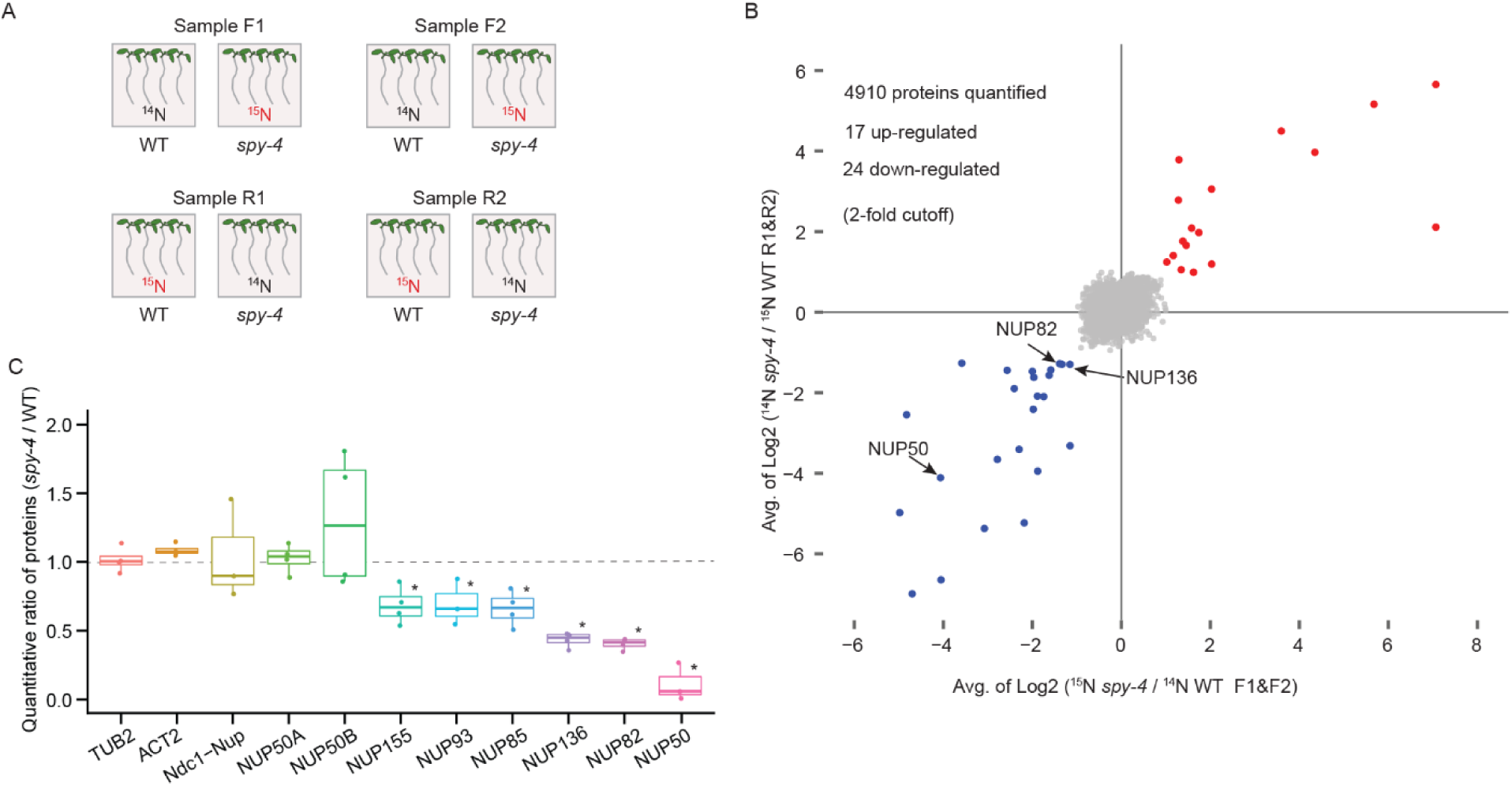
SILIA quantitative proteomics shows that the protein levels of members of the nucleoporin protein family are significantly lower in *spy-4* mutants. (A) Schematic workflow for SILIA quantification of proteomes between WT and *spy-4*. Four biological replicates were performed, including two forward and two reverse labeling experiments. (B) Proteins quantified in at least 3 out of 4 replicates were retained and further filtered as described in Methods. The x-axis is the average of the forward labeling ratio, and the y-axis is the average of the reverse labeling log2 ratio, showing that the nucleoporin proteins NUP50, NUP82, and NUP136(NUP1) are downregulated in *spy-4* mutants using a 2-fold cutoff. (C) Boxplot of nucleoporin protein quantification. Tubulin (TUB2) and actin (ACT2) are used as controls. Asterisks: p<0.01, two-tailed t-test.

This shows that SPY-mediated fucosylation affects the protein level of nucleoproteins. Interestingly, O-GlcNAcylation has been shown to affect both nucleoporin stability (25) and the activity (26), with the sugar modification preventing the NUPs from ubiquitination and degradation and facilitating protein trafficking. Our data provide evidence that O-fucosylation on nucleoporin proteins may play a similar role in regulating their stability.

## Discussion

In this study, we compared and optimized peptide-level and protein-level biotinylation protein enrichment for mapping the signaling network of the low abundance bait SPY, an O-fucose transferase. In total, we enriched 160 interactomes, including 60 known SPY targets and 100 novel potential SPY interactors. This is a significant improvement over our initial IP-MS and PL-MS studies. We also used quantitative proteomics to confirm that SPY affects the protein levels of some of the detected nucleoporin proteins. Our studies highlight PL-MS as a powerful tool for unraveling the complexity of protein interactions.

Protein-level enrichment of biotinylated proteins has been widely adopted due to its relative simplicity in both sample handling and data processing. Here, we present optimized washes to improve PL-MS performance by addressing issues such as collapsed beads sticking to tubes. Our modified washes, which mitigate collapse, minimize sample loss and save time by eliminating the need to scrape beads.

However, peptide-level enrichment has not been well explored, so we sought to optimize the workflow. We have shown that both M280 and MyOne streptavidin beads outperform the antibody-based beads in peptide-level enrichment, possibly due to their higher binding affinity, which allows for extensive washes to remove unmodified peptides. In addition to these types of beads, tamavidin 2-REV beads have been previously validated for peptide-level enrichment (27). This approach uses biotin for elution, but the eluted biotinylated peptides must be further purified to remove contaminating ions that associate with the biotin solution, or the biotin solution must be pre-purified (7). Our procedure does not involve elution with biotin, so we did not observe similar contaminating ions in our peptide enrichment. In addition, we have also shown that a higher concentration of biotin treatment significantly improves the data quality, thereby increasing the confidence and sensitivity in identifying the biotinylated peptides. This feature may also provide a unique advantage for mapping dynamic interactions, such as stimulus-dependent interactions.

Our comparison between MSFragger and Maxquant for LFQ shows that MSFragger has a much better performance for peptide-level enrichment of biotinylated peptides than Maxquant. We show that MSFragger is much more sensitive in detecting the biotinylated peptides than Maxquant, and since both MSFragger and Maxquant are based on detection followed by quantification, better detection results in better quantification. Despite the better performance, we also found that MSFragger still missed some identifications. Currently, none of these software programs consider the use of biotin signature ions, such as m/z 310.16, which we have found to be consistently shown in high intensity in biotinylated spectra. If fragmentation peak lists can be pre-filtered with m/z 310.16, the search may be improved in the future to better identify biotinylated peptides.

The peptide-level enrichment for biotinylated proteins provides a complementary result to the protein-level enrichments. For example, we found that known SPY targets such as ECT5 and unknown interactors such as RAF24, were detected but not shown to be statistically enriched after protein-level enrichment. However, both show statistical enrichment in the peptide-level enrichment.

With all the improvements, we have identified 60 known targets and 100 new SPY interactors. Some of the new interactors belong to the same family members as the known targets, suggesting that they may also be SPY targets. Other interactors may have a regulatory role for SPY. Notably, several proximity labeling experiments have been used to identify sugar-modified targets as well as regulators of sugar cycle enzymes, for example, OGA-BioID was used to probe oxidative stress-induced binding proteins to OGA (28), or the TPR domain of human OGT (HsOGT) was fused to probe the HsOGT interactome (29). In addition, a sugar-binding domain was fused to TurboID to allow spatiotemporal tracking of GlcNAcylated “activity hubs” during cellular signaling (30).

Despite the many improvements, we are aware that there are still false negatives. We believe that there are two main sources: high background and low signal-to-noise of biotinylated interactors. We have observed a high background similar to that observed in (4). We note that the streptavidin beads were hydrophobic beads, leading us to speculate that the bFexpreseads may attract proteins with hydrophobic domains, resulting in a high background. While it is difficult to test this at the protein level, we examined the behavior of the peptides after enrichment at the peptide level. Indeed, we observed that a significant amount of hydrophobic unmodified peptides was retained even after extensive washes (Supplementary Fig. S2). On the other hand, some target proteins may be randomly biotinylated by the TD-YFP control, so that their signal-to-noise would be low, resulting in a false negative, as shown by the yellow dots (SPY targets) overlaid with grey dots (not significantly enriched SPY-TD/TD-YFP) in Supplementary Fig. S6A and B (Supplementary Table 7).

Our improved methods will be used not only for TurboID approaches, but also for other PL-MS, such as APEX, and not only for PPI, but also for other subcellular proteomics as well as cell type proteome mapping. In particular, the enhancement of peptide-level enrichment can help provide topological information for subcellular proteomics. In the future, we can take advantage of signature ions such as imoKbiotin (310.16), for example, using the signature ion to trigger MS3 acquisition in TMT datasets, or pre-filtering the peak list with the signature ion, and integrating the signature ions into the data search score algorithm to improve the detection of biotinylated peptides.

## DATA AVAILABILITY

The mass spectrometry proteomics data have been deposited to the ProteomeXchange Consortium via the PRIDE partner repository. All other data are available from the corresponding author on reasonable request. The spectra for the biotinylated peptide identification can be viewed using MSviewer (31).

## Acknowledgments

We would like to thank Robert Chalkley and Peter Baker for providing technical expertise. We would also like to thank Prof. Neil Olszewski for sharing the *spy*-4 mutants. We thank Andrea Mair and Prof. Dominique Bergmann for sharing the TurboID constructs. This work was funded by the National Institutes of Health grants R01GM135706 and S10OD030441 to S.-L.X. and by the Carnegie Endowment Fund to the Carnegie Mass Spectrometry Facility.

## Author contributions

S.-L.X. conceptualization; T.G, S.K. Biotinylated protein and peptide-level experiments; R.S., WT/*spy-4* SILIA proteome comparison. T.G., S.K, R.S., A.V.R. data analysis; S.K., D.B, constructs and genetic analysis. W.N. IP-MS analysis. T.G., S.K Figures. T.G., S.K., S.-L.X. manuscript writing.

## Methods

### Construct Generation and Transformation

The pENTR1A-SPY construct was generated by amplifying the full-length coding sequence of SPY using PCR, which was then cloned into the pENTR1A vector (Thermo Fisher Scientific). Approximately 2 kb upstream of the SPY ATG start codon, which encompasses the promoter and 5’-untranslated leader (32, 33) was amplified and cloned into the pENTR5 vector gateway clone to generate the SPY promoter entry clone (pENTR5-SPYpro). Three entry vectors (pENTR5-SPYpro, pENTR1A-SPY, pDONR_P2R-P3_R2-Turbo-mVenus-STOP-L3) were recombined with destination vector pB7m34GW (4) to generate the final expression vector SPYpro::SPY-TD-mVENUS (abbreviated as SPY-TD). The control TurboID-YFP entry vector (pENTR_L1-Turbo-YFP-STOP-L2) was made by applying site-directed mutagenesis to introduce a stop codon right before the NLS of the entry clone (pENTR_L1-Turbo-YFP-NLS-STOP-L2) (4) to eliminate the NLS signal sequence. The control construct SPYpro::TD-YFP (abbreviated as TD-YFP) was generating by recombining two entry vectors (pENTR5-SPYpro, pENTR_L1-Turbo-YFP-STOP-L2) with destination vector R4pGWB501 to generate the final expression vector. SPY-TD was transformed into the *spy-3* mutant and TD-YFP transformed into wild type via the floral-dip method (34) and selected based on BASTA and hygromycin, respectively. Complemented SPY-TD lines and TD-YFP lines with a similar expression level to that of SPY-TD were chosen for further analysis.

### Complementation phenotype examination

PAC resistance. Wild-type, *spy* mutant, and three independent transgenic lines were germinated in the presence of the gibberellin biosynthesis inhibitor, paclobutrazol (PAC). Sterilized seeds were plated on ½ Murashige & Skoog (MS) media (2.17g/L MS, 6 g/L Phytoblend, pH 5.8) (control) and ½ MS medium supplemented with 15 μg/ml PAC (treatment). The seeds were placed in 4 °C cold room for 2 days for stratification then transferred to a chamber at 22 °C with a 24h-light (light intensity of 82 μmol m−2 s−1) for 10 days. Germination in the presence of PAC was assessed in the transgenic lines compared to wild type and *spy-3* mutant.

Leaf Serration. Phenotypic analysis of SPY-TD transgenic lines was performed based on leaf serration (presence or absence of serration). Leaves were assessed from three weeks old plants grown in greenhouse conditions with 100 µmol light intensity.

### Confocal Microscopy

Confocal microscopy images of Arabidopsis seedlings expressing TurboID constructs were taken with a Leica SP8 microscope. Images were processed using Adobe photoshop software.

### Plant Growth and Biotin Treatment

The SPY-TD and TD-YFP transgenic seedlings were sown on ½ MS medium, cold-treated for 3 days, then placed vertically in a growth chamber under 24hr light conditions at 22 °C for 10 days. 10-day seedling tissues (1 or 2 g, three replicates per condition) were harvested and submerged in 30 ml of biotin solution (50-500 μM) for 1 hour at room temperature. Following treatment, seedlings were washed thoroughly with deionized water to remove excess biotin, patted dry, and flash frozen in liquid nitrogen. Frozen tissue was ground to a fine powder using CryoMill and stored at −80°C until protein extraction.

### Streptavidin Immunoblot

For the immunoblot, 10-day-old Arabidopsis seedlings were treated with various concentrations of biotin (0-1000 µM) for 1 hour, and 40 mg of fresh tissues from each sample were flash-frozen after removing excess biotin. Total proteins were extracted into three volumes of 2X SDS sample loading buffer, and proteins were separated by SDS-PAGE and blotted onto a PVDF membrane (0.2 µm, BioRad). After blocking with 3% BSA, the blot was probed with anti-streptavidin antibody (1:2000, Thermo Fisher Scientific).

### Enrichment of Biotinylated proteins at the Peptide-level

#### Initial Bead type comparison

Protein extractions were performed using phenol extraction and digested to peptides as described in (35). Dried peptides were resuspended in 1X PBS buffer, then divided into three equal fractions for three beads type comparison. Each fraction (∼4.5 mg peptides) was used for peptide enrichment with either M280 streptavidin beads (Invitrogen), MyOne streptavidin beads (Invitrogen), or anti-biotin bound agarose beads (Cell Signaling Technology).

Anti-biotin beads enrichment was performed similarly as described in (8), following the manufacturer’s suggestions with modifications as follows. Antibody bead slurries were washed with 1mL of 1X PBS buffer four times. About 4.5 mg Peptides were resuspended in 1 ml 1X PBS buffer and incubated with anti-biotin agarose beads (80 μL of bead slurry, neat bead volume ∼ 60 μL) for 2 hours at 4 °C with gentle rotation. Beads were washed twice with 1X PBS buffer and three times with HPLC-grade H_2_O. Peptides were eluted with a total of 105 μL of elution buffer (80% ACN, 0.2% TFA), dried and desalted.

Similarly, peptides resuspended in 1 ml of 1X PBS buffer were incubated with MyOne or M280 streptavidin beads (100 μL bead slurry, prewashed with PBS buffer, neat volume <10 μL) for 1 hour at room temperature on a rotator, washed with 1X PBS buffer twice, 1X PBS with 5% ACN four times, and 5% ACN in ddH_2_O once. Biotinylated peptides were eluted with 600 μl pre-heated elution buffer (0.2% TFA, 0.1% FA, and 80% ACN) at 75°C. Eluted peptides were dried, resuspended in 5% ACN and 0.1% TFA, then desalted.

#### Testing acetonitrile (ACN) concentrations in wash buffers

Biotinylated peptide enrichment was evaluated using 5% ACN, 10% ACN, and 15% ACN washes. M280 streptavidin beads (Invitrogen) were used for the enrichment process. Dried peptides (∼9 mg) were resuspended in 1 mL of 1X PBS with the designated ACN percentage and introduced to pre-washed 100 μL of M280 beads (washed with 200 μL 1X PBS with the respective ACN percentage). The peptide-bead mixture was incubated for 1 hour at room temperature on a rotor and washed four times with 1 mL of 1X PBS with the designated ACN percentage, followed by a final wash with 1 mL of the respective ACN percentage in HPLC grade H_2_O. Peptides were eluted as mentioned above.

#### Testing the effects of increased biotin concentration for biotin treatment

Harvested plant tissues were biotin treated as described previously using 500 μM biotin solution. All subsequent freezing and grinding steps are the same. Biotinylated peptide enrichment with 100 μL of M280 streptavidin bead slurry was performed as follows: peptides were resuspended in 1 mL of 1xPBS in 15% ACN, binding at 1 hour at room temperature, and washed with PBS in 15% ACN four times, and 15% ACN in HPLC grade H_2_O twice. Peptides were eluted, dried, and desalted as mentioned above.

### Biotinylated protein enrichment at the protein-level

Biotinylated protein enrichment at the protein-level was performed the same as described in (3, 4), except the wash steps. Briefly, the proteins were extracted in extraction buffer (50 mM Tris pH 7.5, 150 mM NaCl, 0.1% SDS, 1% NP-40, 0.5% Na-deoxycholate, 1 mM EGTA, 1 mM DTT, 1x complete protease inhibitor, 1 mM PMSF), desalted using PD-10, and incubated with 200 µL pre-washed MyOne Streptavidin C1 Dynabeads (Invitrogen) on a rotor wheel at 4°C overnight, followed by extensive washes. Samples were divided into two equal fractions. One set of samples was treated with standard wash methods as described in (3, 4), including twice with cold extraction buffer (beads were then transferred into a new tube), twice with cold equilibration buffer (50 mM Tris pH 7.5, 150 mM NaCl, 0.1% SDS, 1% NP-40, 0.5% Na-deoxycholate, 1 mM EGTA, 1 mM DTT), once with cold 1 M KCl, twice with cold 100mM Na_2_CO_3_, and once with 2M urea in 10 mM Tris pH 8 at room temperature. The second set of samples was treated with a modified wash method in which the Na_2_CO_3_ and urea washes are replaced with extraction buffer washes.

Bead samples of both standard and modified washes were spun down to remove the remaining wash buffer, then washed twice with 1 mL of 50 mM Tris pH 7.5, then with 1 M urea in 50 mM Tris pH 7.5. Wash buffer was removed and replaced with 80 µL of trypsin buffer (50mM Tris pH 7.5, 1M Urea, 1 mM DTT, 0.4 µg Trypsin). Beads were incubated at 25°C for 3 hrs, after which the supernatant (containing peptides) was transferred to a new tube. More supernatant was collected after two additional washes with 60 µl of 1M urea in 50 mM Tris pH 7.5 and combined with the original supernatant resulting in approximately 200 uL of eluted peptides.

Iodoacetamide was added to the sample at a final concentration of 10mM and incubated for another 45 mins at 25°C. Trypsin (0.5 µg) was introduced to the samples for overnight digestion. The following morning an additional 0.5 µg of trypsin was added and allowed to digest for 4 more hours. The reaction was quenched by adding formic acid (FA) to a final concentration of 1%. Samples were centrifuged for 20 mins, desalted using C18 P-10 Agilent ZipTips, and dried. Each sample was resuspended in 0.1% FA directly prior to MS injection.

### Immunoprecipitation of the SPY Interactomes (IP-MS)

The SPY-GFP (13) and TAP-GFP seedlings (36) were grown for 7 days at 21°C under constant light on ½ MS medium. Tissues were harvested, ground in liquid nitrogen, and stored at −80°C. Immunoprecipitation was performed as described previously with slight modifications. Briefly, proteins were extracted in MOPS buffer (100 mmol/L MOPS, pH 7.6, 150 mmol/L NaCl, 1% (v/v) Triton X-100, 1 mmol/L PMSF, 2× Complete protease inhibitor cocktail, and PhosStop cocktail (Roche)), centrifuged, and filtered through two layers of Miracloth, followed by incubation with a modified version of LaG16-LaG2 anti-GFP nanobody conjugated to Dynabeads (Invitrogen) for 3 h at 4°C. Incubation was followed by four 2-minute washes with immunoprecipitation buffer and eluted with 2% (w/v) SDS buffer containing 10 mmol/L tris(2-carboxyethyl) phosphine (TCEP) and 40 mmol/L chloroacetamide at 95°C for 5 minutes. Eluted proteins were separated by SDS-PAGE. After Colloidal Blue staining, the whole lane of protein samples was excised in three segments and subjected to in-gel digestion with trypsin. Three biological experiments were performed.

### SILIA quantitative proteome analysis between WT and *spy-4* mutant

The SILIA proteome quantification between the WT and *spy-4* plants were performed as described in (18). Briefly, WT and *spy-4* were grown for 14 days on Hoagland’s medium containing ^14^N or ^15^N (1.34 g/L Hoagland’s No. 2 salt mixture without nitrogen, 6 g/L Phytoblend, and 1 g/L KNO_3_ or 1 g/L K^15^NO_3_ (Cambridge Isotope Laboratories), pH 5.8, 1% sucrose). Plates were placed vertically in a growth chamber under constant light condition at 21– 22 °C. Whole plant tissues were harvested in liquid nitrogen. Proteins were extracted from eight samples (two ^14^N-labeled WT samples – 1 and 5; two of ^15^N-labeled WT samples – 2 and 6; two of ^14^N-labeled *spy-4* samples – 3 and 7; and two ^15^N-labeled *spy-4* samples – 4 and 8) individually using 2X SDS sample buffer (plant tissue mass (mg): Buffer (μL) ratio = 1:3) and mixed as follows: two forward samples F1 and F2 (^14^N WT/^15^N *spy-4*, Mix1+4; Mix 5+8) and two reverse samples R1 and R2 (^14^N *spy-4*/^15^N WT, Mix 2+3; Mix 6+7) and separated by the SDS-PAGE. Five segments were excised, and trypsinized. The peptide mixtures were desalted using C18 ZipTips (Millipore) and analyzed by liquid chromatography-mass spectrometry (LC-MS).

### SILIA WT vs *spy-4* data analysis/filtering

SILIA data containing two forward replicate experiments (F1, F2) and two reverse labeling replicates experiments (R1,R2) were searched and quantified the same as described by (18). SILIA ^14^N /^15^N labeled WT and *spy-4* data were filtered to remove poor quality identifications. Because ^15^N quantification can suffer from contamination that can interfere with quantification, data was further filtered using the following criteria: 1) Proteins quantified in three or more replicates were retained. 2) Proteins with log2-fold enrichment values between −1 and 1 were retained. 3) Proteins with all values either greater than or equal to 1 or less than or equal to −1 (for up- or down-regulated proteins) were retained. After filtering, a total of 4910 proteins are quantified.

### LC-MS/MS analysis

Peptides were analyzed by liquid chromatography–tandem mass spectrometry (LC-MS) on an EasyLC1200 system (Thermo) connected to a high-performance quadrupole Orbitrap mass spectrometer Eclipse (Thermo). Peptides were first trapped using trapping column Acclaim PepMap 100 (75 µm x 2cm, nanoViper 2Pk, C18, 3 µm, 100A), then separated using analytical column AUR2-25075C18A, 25CM Aurora Series Emitter Column (25cm x75 µm, C18, 1.6um) (IonOpticks). The flow rate was 300 nL/min. For peptide-level enrichment samples, peptides were eluted by a gradient from 3 to 10% solvent B (80% (v/v) acetonitrile/0.1% (v/v) formic acid) over 1 min, 10 to 35% solvent B over 105 min, from 35 to 44% solvent B over 15 min, followed by a short wash for 15 min by 90% solvent B. For protein-level enriched samples, peptides were eluted by a gradient from 3 to 5% solvent B over 1 min, 5 to 28% solvent B over 105 min, from 28 to 44% solvent B over 15 min, followed by a short wash (15 min) at 90% solvent B.

For data acquisition, Precursor scan was from mass-to-charge ratio (m/z) 375 to 1600 (resolution 120,000; AGC 200,000, maximum injection time 50ms, Normalized AGC target 50%, RF lens(%) 30) and the most intense multiply charged precursors were selected for fragmentation (resolution 15,000, AGC 5E4, maximum injection time 22ms, isolation window 1.4 m/z, normalized AGC target 100%, include charge state=2-8, cycle time 3 s). Peptides were fragmented with higher-energy collision dissociation (HCD) with normalized collision energy (NCE) 27. Dynamic exclusion was enabled for 30s.

For IP-MS and SILIA samples, Peptides were analyzed by liquid chromatography–tandem mass spectrometry (LC-MS) on an EasyLC1200 system (Thermo) connected to a high-performance quadrupole Orbitrap Q Exactive (Thermo). Peptides were separated using ES803 (50cm) column. The LC gradient is the same as above. Precursor scan was from mass-to-charge ratio (m/z) 375 to 1600 (resolution 120,000; AGC 3.0E6, maximum injection time 100 ms) and top 20 most intense multiply charged precursors were selected for fragmentation (resolution 15,000, AGC 5E4, maximum injection time 60ms, isolation window 1.0 m/z, minimum AGC target 1.2e3, intensity threshold 2.0 e4, include charge state =2-8). Peptides were fragmented with higher-energy collision dissociation (HCD) with normalized collision energy (NCE) 27. Dynamic exclusion was enabled for 24s.

### Search with Protein Prospector, Maxquant, MSFragger

For Protein Prospector (version 6.4.9), MS/MS data was converted to peaklists using a script PAVA (peaklist generator that provides centroid MS2 peaklist (18). Raw MS/MS data was loaded in Maxquant (v2.2.0 for peptide-level enrichment and v2.4.12 for protein-level enrichment) or MSFragger (v20.0) for peptide identifications and LFQ.

In all three programs data was searched against the TAIR10 database *Arabidopsis thaliana* (https://www.arabidopsis.org/), concatenated with decoy protein sequences (a total of 35386 entries). A 10 ppm precursor mass tolerance and 20 ppm MS/MS2 tolerance was allowed. Carbamidomethyl (cysteine (C)) was searched as a constant modification. For peptide-level enriched samples, variable modifications allowed include biotinylation at uncleaved lysines (Biotin(K)), protein N-terminal acetylation, methionine (M) oxidation, N-terminal acetylation with M oxidation, N-terminal M-loss, N-terminal M-loss with acetylation, and peptide N-terminal Gln conversion to pyroglutamate, with three maximum variable modifications per peptide. Cleavage specificity was set to trypsin, allowing two missed cleavages. For protein-level enriched samples, all the above listed variable modifications were included except biotin modification. Two maximum modifications and one missed cleavage per peptide were allowed. For all samples, a 5% FDR was permitted for protein and 1% FDR permitted for peptide identifications.

For peptide-level enrichment with Maxquant, LFQ was run with min ratio count as 1, match time window 0.7 min, and other parameters as default. For protein-level enrichment samples, Maxquant LFQ was run with min ratio count as 2 and match time window with 0.7 min, and other parameters as default. For MSFragger, the LFQ was run on default settings. Match between runs was used for both Maxquant and MSFragger searches.

### Statistics processing using Perseus

For peptide-level enrichment either Maxquant’s ‘peptides.txt’ file or MSFraggers’s ‘combined_modified_peptides’ was loaded into Perseus v2.0.10, with LFQ intensities specified as ‘Main’ columns. Identifications were filtered to only include biotinylated peptides for further analysis. For protein-level enrichment either Maxquant’s ‘proteinGroups.txt’ file or MSFragger’s ‘combined_proteins’ was input to Perseus, also with LFQ intensities specified as ‘Main’ columns. Decoys and/or contaminants were removed, then the LFQ intensities were log2 transformed and quality checked with histograms. Non-reproducible identifications were removed (keep only rows with at least 3 valid values in either SPY-TD replicates or TD-YFP replicates), then imputation from a normal distribution was performed to replace remaining missing values. A two-sided T-test with S0=0.5 and FDR=0.05 was used to determine significance and enrichment of each identified peptide or protein.

### SILIA data Search and quantification

MS/MS data were converted to peaklist using a script PAVA (peaklist generator that provides centroid MS2 peaklist), and data were searched using Protein Prospector using the same TAIR database. A precursor mass tolerance was set to 10 ppm and MS/MS2 tolerance was set to 20 ppm., Carbamidomethyl (C) was searched as a constant modification. Variable modifications include protein N-terminal acetylation, peptide N-terminal Gln conversion to pyroglutamate, Met oxidation. The search and quantification were the same as described by (18).

---

**Supplemental Figure S1.**
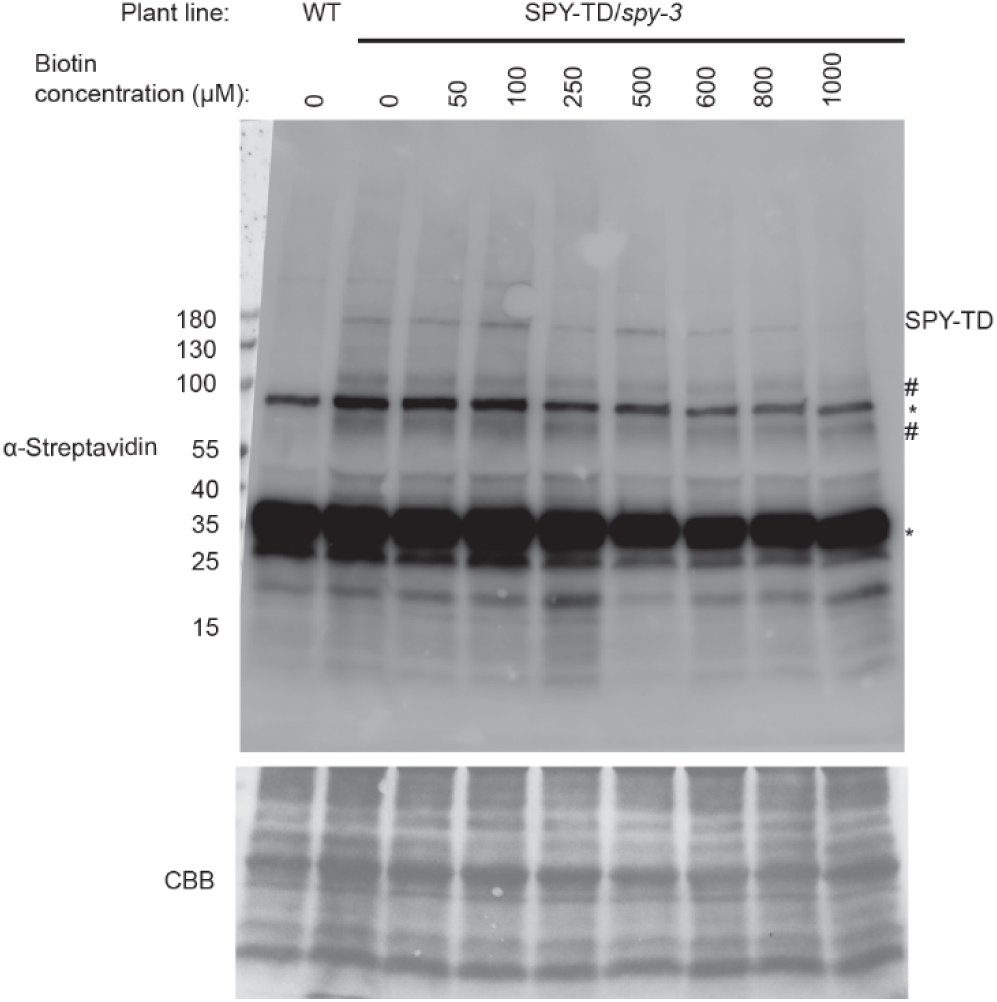
SPY-TD shows no visible increase in biotinylation activity with increasing biotin concentration using anti-streptavidin western blotting. Ten-day-old seedlings of SPY-TD lines were treated with biotin at concentrations ranging from 0-1000 µM for 1 hour. Immunoblots using streptavidin-HRP show the activity of the SPY-TD (top panel). Coomassie Brilliant Blue (CBB) staining is used as a loading control (bottom panel). Asterisks (*) mark endogenous biotinylated protein positions. Each sample was taken from a pool of 40 mg tissue. The SPY-TD band and two biotinylated bands in the highlights are marked with #.

**Supplementary Figure S2.**
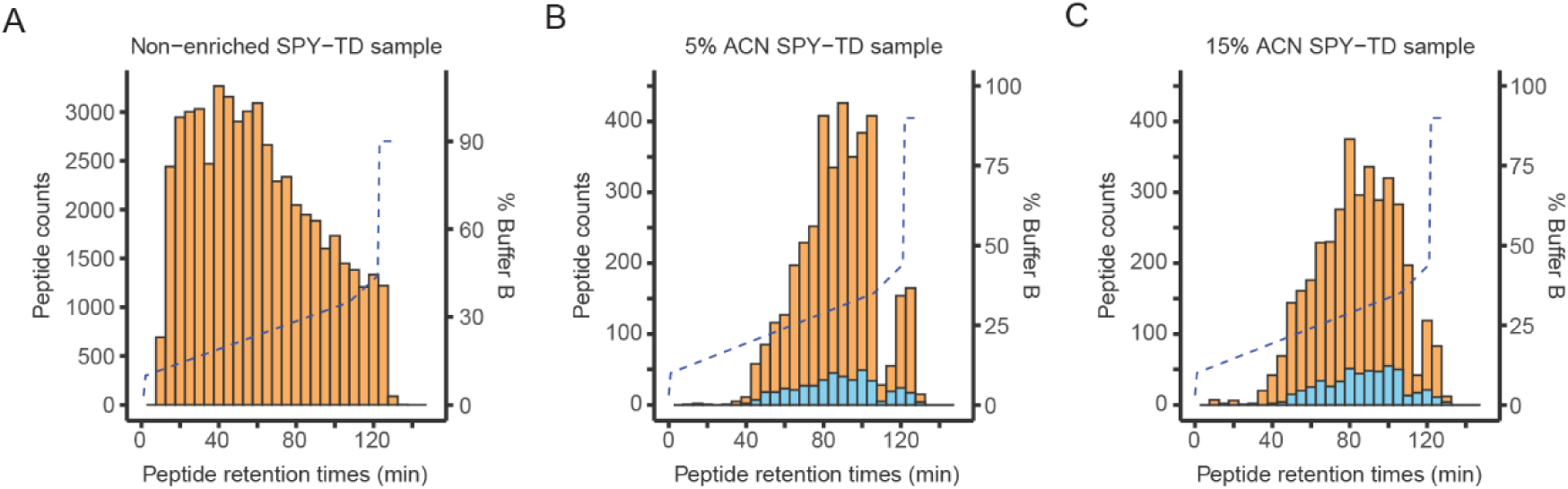
Biotinylated peptides show higher hydrophobicity and unmodified hydrophobic peptides were retained by M280 streptavidin beads. Peptide distributions across gradient runs are shown for non-enriched and enriched peptide samples. The x-axis represents retention time (RT) and y-axis the number of peptides. The gradient is shown by the dashed line as the buffer B (80%ACN, 0.1%FA) increases over time. Each histogram bar represents the number of peptides eluted within a 5-minute segment. (A) RT distribution of peptides from an unenriched SPY-TD sample. (B-C) RT distribution of enriched peptides from SPY-TD samples treated with 5% ACN or 15% ACN washes. Biotinylated peptides are superimposed in blue color.

**Supplementary Figure S3.**
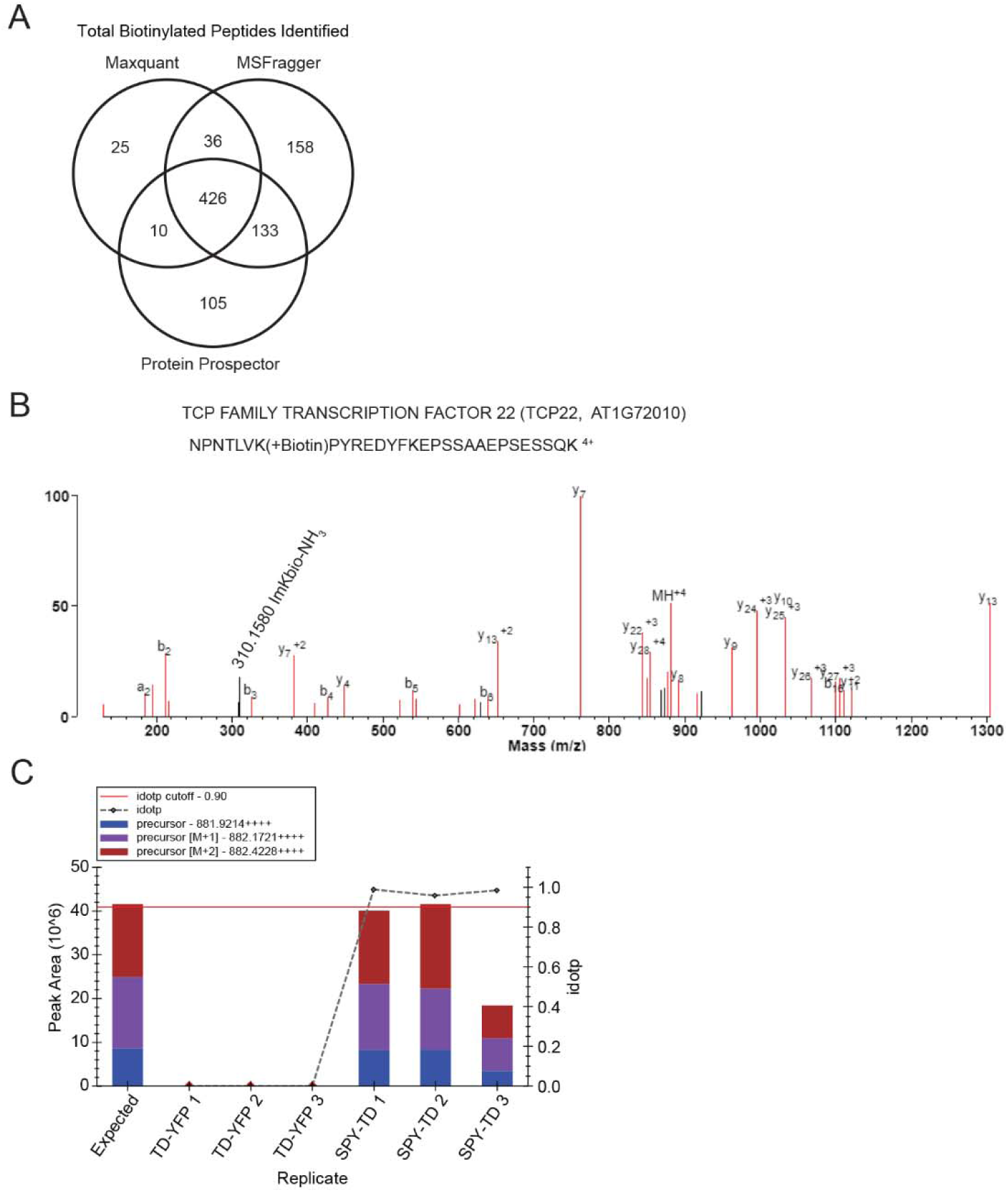
MSFragger performs best in identifying biotinylated peptides compared to Protein Prospector and Maxquant but may still miss some identifications. (A) Venn diagram of total identified biotinylated peptides from SPY-TD and control samples using Maxquant, MSFragger, and Protein Prospector. (B) A highly scored quadruple charged biotinylated peptide (NPNTLVK(+Biotin)PYREDYFKEPSSAAEPSESSQK) from TCP22 is identified by Protein Prospector but not by MSFragger. Signature ion of m/z 310.158 corresponding to the immonium ion of biotinylated lysine with loss of ammonia is labeled. (C) Skyline quantification shows that the TCP22 biotinylated peptide is specifically enriched in the SPY-TD but not in the TD-YFP control.

**Supplementary Figure S4.**
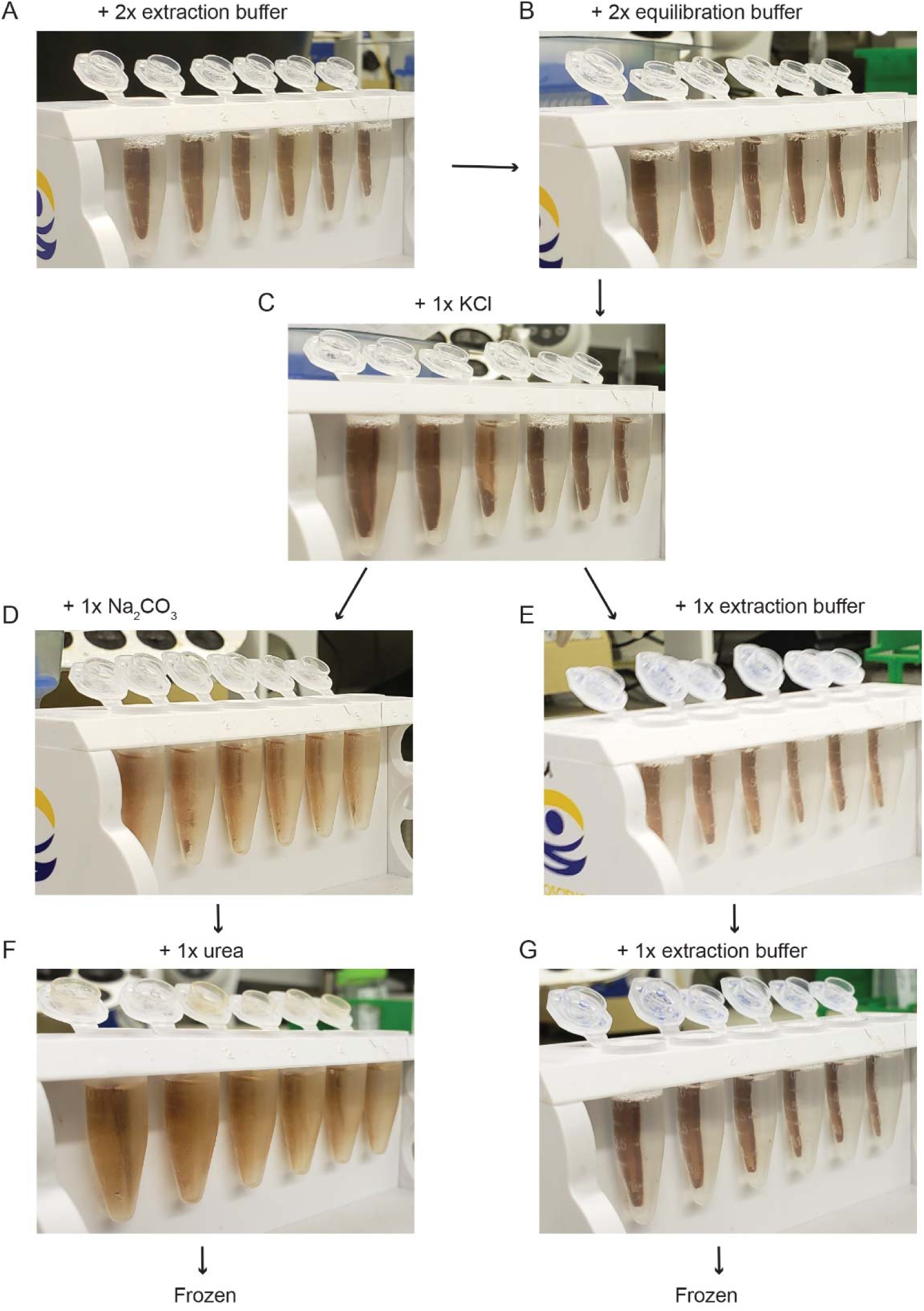
Stepwise comparison of beads between standard and modified washes. (A) Beads on a magnetic rack after two washes of extraction buffer. (B) Beads after two washes of the equilibration buffer. (C) Beads after one KCl wash. After KCl washes, the beads were split, half washed with standard washes (D, F) and half washed with modified washes (E, G) before freezing at −80°C.

**Supplementary Figure S5.**
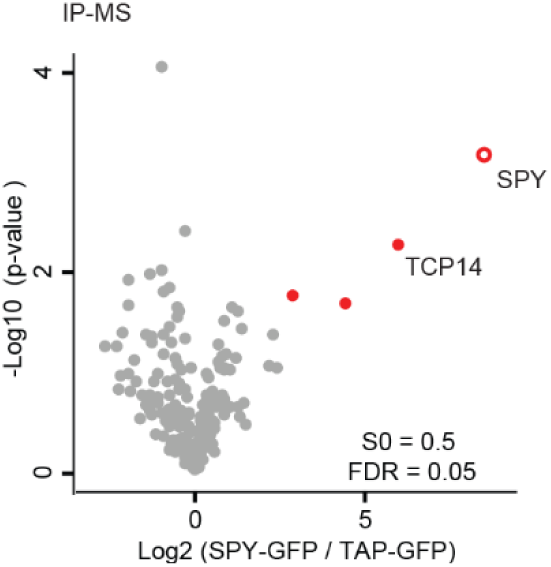
Immunoprecipitation followed by mass spectrometry of SPY-GFP shows specific enrichment of SPY protein, TCP14, and two other proteins. Immunoprecipitation was performed using SPY-GFP and TAP-GFP lines, followed by mass spectrometry analysis and label-free quantification.

**Supplementary Figure S6.**
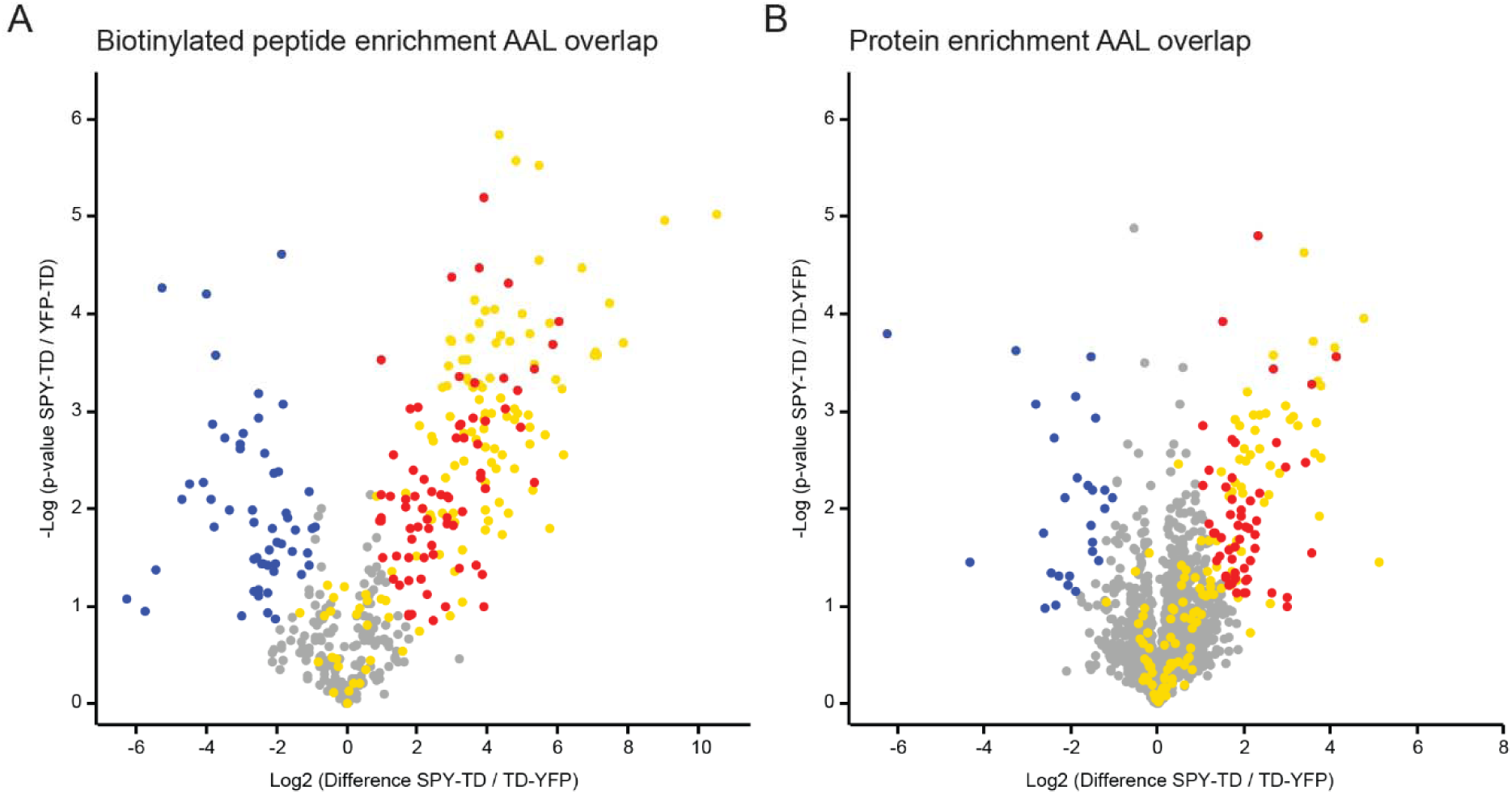
Some known SPY targets are identified but not enriched in SPY-TD samples compared to TD-YFP. All SPY targets enriched by AAL are shown in yellow overlaid on peptide- and protein-level enrichment data. Yellow points overlaid on gray points represent SPY targets enriched by AAL but not significantly different between SPY-TD and TD-YFP. (A) 17 known SPY targets are unenriched in peptide-level enrichment samples. (B) 79 known SPY targets are unenriched in protein-level enrichment samples. 9 known SPY targets that are unenriched in protein-level samples are enriched in peptide-level samples. 74 known SPY targets aren’t enriched in either peptide- or protein-level samples.

## Notes

### Competing Interest Statement

The authors have declared no competing interest.

